# Bioremediation of industrial pollutants by insects expressing a fungal laccase

**DOI:** 10.1101/2021.04.28.441736

**Authors:** Michael Clark, Kate Tepper, Kerstin Petroll, Sheemal Kumar, Anwar Sunna, Maciej Maselko

## Abstract

Inadequate management of household and industrial wastes pose major challenges to human and environmental health. Advances in synthetic biology may help address these challenges by engineering biological systems to perform new functions such as biomanufacturing of high-value compounds from low-value waste streams and bioremediation of industrial pollutants. The current emphasis on microbial systems for biomanufacturing, which often require highly pre-processed inputs and sophisticated infrastructure, is not feasible for many waste streams. Concerns about transgene biocontainment have limited the release of engineered microbes or plants for bioremediation. Engineering animals may provide opportunities for utilizing various waste streams that are not suitable for microbial biomanufacturing while effective transgene biocontainment options should enable *in situ* bioremediation. Here, we engineer the model insect *Drosophila melanogaster* to express a functional laccase from the fungus *Trametes trogii*. Laccase expressing flies reduced concentrations of the endocrine disruptor bisphenol A by more than 50% when present in their growth media. A lyophilized powder made from engineered adult flies retained substantial enzymatic activity, degrading more than 90% of bisphenol A and the textile dye indigo carmine in aqueous solutions. Our results demonstrate that transgenic animals may be used to bioremediate environmental contaminants *in vivo* and serve as novel production platforms for industrial enzymes. These results support further development of insects, and possibly other animals, as bioproduction platforms and their potential use in bioremediation.

## Main Text

Developing economically viable approaches for sustainable waste management and mitigation of environmental pollution are important for ensuring a high quality of life^1^. At present, over half of all municipal solid waste is either landfilled^2^, where it is estimated to contribute to 11% of global methane emissions^3,4^, or openly dumped, where it contaminates drinking water and disrupts natural ecosystems^5^. Pollution from industrial wastes includes a variety of persistent compounds which have been linked to neurological defects, cancer, and species extinction^6–10^.

Regulatory changes including prohibiting discharge of industrial pollutants and reducing the landfilling of food waste are helpful, however, the cost of compliance can be substantial^11^. Generating value from a variety of waste-streams and developing inexpensive methods to treat pollutants *in situ* will reduce the costs of achieving sustainability targets and help support a circular economy^12^.

We propose developing solutions by expanding the enzymatic repertoire of select animals to consume waste, degrade xenobiotic chemicals, and produce high-value compounds (Fig. 1). Microbial and fungal enzymes can enable animals to breakdown pollutants *in situ*. These may include herbivores that degrade pollutants accumulating in plants, fish that secrete enzymes to degrade aquatic pollutants, or invertebrates engineered to interrupt the biomagnification of xenobiotics in food webs. Effective transgene biocontainment safeguards are currently a challenge for plants/microbes^13,14^, but are possible in animals via surgical sterilization or engineered genetic incompatibilities^15^. Developing insects as a biomanufacturing platforms for industrial enzymes, lipids, or other molecules is also an unexplored avenue for waste valorisation.

**Fig. 1:**
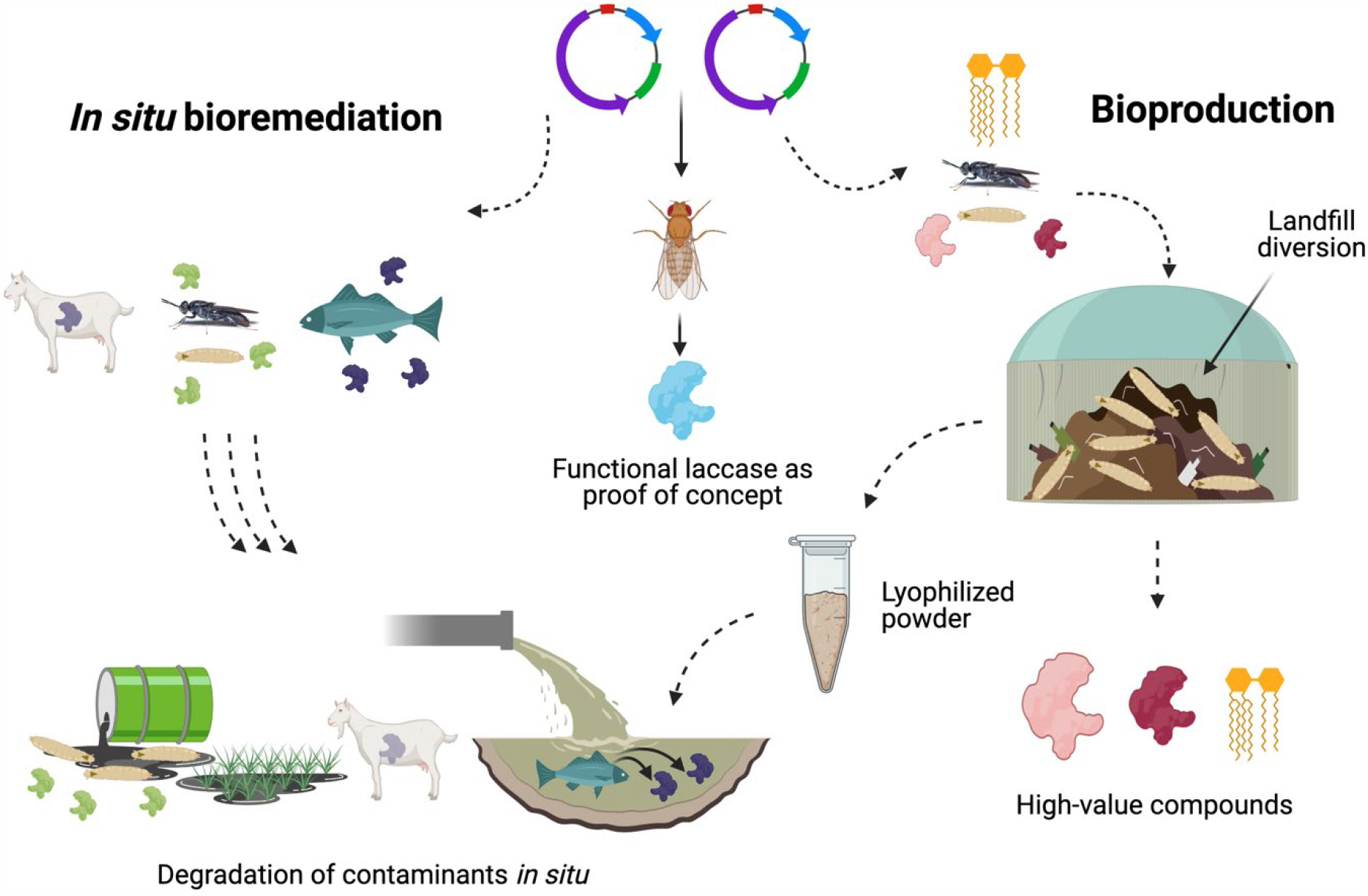
Applications of transgenic animals for bioremediation and bioproduction.

Insects have advantages over bacteria and fungi for processing many waste-streams. Their complex digestive systems, including mastication, allow them to consume a variety of low-value wastes, including municipal organic waste^16^ and livestock manure^17^. Crude waste can be fed directly to insects without pre-processing, refinement, and sterilization necessary for microbial fermentation^18,19^. Some insect species, such as Black soldier flies (*Hermetia illucens*), which are already used to manage waste have excellent tolerance to bacteria and fungi^20^ and are easily separated from the organic waste for downstream processing^21^. Insect biotechnology is readily scalable for mass production, as the infrastructure can be relatively unsophisticated, and the installation costs and land use requirements are low^22^. Furthermore, utilising organic waste means crops that could otherwise be used for food do not need to be diverted as feedstocks for fermentation, and will not threaten food security.

There is a long history of using lepidopteran insect larvae to make recombinant proteins starting with *Bombyx mori* expressing human α-interferon^23^. However, these utilize transient baculoviral systems to infect the larvae. This approach is not practical for large-scale or long-term, multi-generational use. Here we explore the potential of stably engineered insects as platforms for the production of an industrially relevant fungal laccase and bioremediation of polluted sites. Laccases are multicopper oxidases that are present in many taxa. Fungal laccases were selected due to their demonstrated utility for bioremediation of a wide variety of industrial pollutants such as bisphenols^24^, industrial dyes^25^, pharmaceuticals^26^, phenolics^27^, polycyclic aromatic hydrocarbons^28^, perfluoroalkyl and polyfluoroalkyl substances (PFAS)^29,30^, plastics^31^, pesticides^32^, and mycotoxin contaminants such as aflatoxins^33^. Fungal laccases also have diverse industrial applications in the textile, paper and pulp, food, pharmaceutical, chemical synthesis, and forestry industries^34–36^. They have been applied to treat distillery^37^, paper and pulp^38^, and olive mill^39^ industry wastewater. Ligninolytic enzymes, such as fungal laccases, also show potential application in the delignification of feedstocks for biofuel production^40^. However, fungal laccases have not yet reached appreciable commercial scale production. This is because the native hosts for laccases often produce low yields and although this has been improved using heterologous hosts, the production levels are still too low for many commercial applications^41^.

We engineered the common fruit fly (*Drosophila melanogaster*) to express a laccase from the fungus *Trametes trogii*. Transgenic flies degraded bisphenol A (BPA), a widespread endocrine disruptor^42^, when present in their diet. We also demonstrate that aqueous solutions of BPA and indigo carmine (IC), a textile dye and aquatic pollutant^43^, can be treated with a lyophilized powder from adult transgenic flies to degrade both compounds. Our results describe the first instance of engineering an animal to perform an ecosystem service by heterologous expression of a fungal enzyme and demonstrate the potential for using transgenic strains of insects for enzyme bioproduction.

## Results

### Screening for fungal laccase activity in transgenic *D. melanogaster*

*D. melanogaster* and other insects express endogenous laccases involved in immune defense^44^. However, it was unknown if an insect could heterologously express a functional fungal laccase or if its expression would be toxic. We were also uncertain which tissues would provide the optimal expression of a functional laccase for either *in vitro* or *in vivo* activity. We selected a short tubulin promoter, previously used to express a toxic gene^45^, with the goal of achieving moderate expression across many tissues. The native fungal signal peptides were replaced with the *D. melanogaster* larval cuticle protein 9^46^ signal peptide to facilitate extracellular secretion.

We generated homozygous transgenic strains expressing laccases from two fungal species, *T. trogii* and *Polyporus brumalis*, known to have a broad substrate range and high redox potential (Table 1). Fly lysates from both were assayed for oxidation of the chromogenic laccase substrate 2,2’-Azino-bis(3-ethylbenzthiazoline-6-sulfonic acid) (ABTS). The engineered *D. melanogaster* strain Dm/Tt.Lcc1, which expresses a laccase from *T. trogii*, oxidized ABTS (Fig. 2a). There was no activity observed in the wild-type or from the engineered Dm/Pb.Lac1 strain expressing a laccase from *P. brumalis* (Fig. 2a). Moving forward, we focused on characterizing Dm/Tt.Lcc1.

**Table 1.** The fungal laccases engineered into *D. melanogaster*.

| Gene name | Native host | Genbank accession | Redox potential (V vs. NHE, pH 5) | Laccase substrates |
| --- | --- | --- | --- | --- |
| *Lcc1* | *Trametes trogii* | AJ294820.1 | $E^\circ$ 0.79 V <sup>49</sup> | Aminophenols, polyphenols, industrial dyes<sup>50–52</sup>. |
| *Lac1* | *Polyporus brumalis* | ABN13591.1 | $E^\circ$ 0.80 V <sup>53</sup> | Industrial dyes, aromatic hydrocarbons <sup>53</sup>. |

**Fig. 2:**
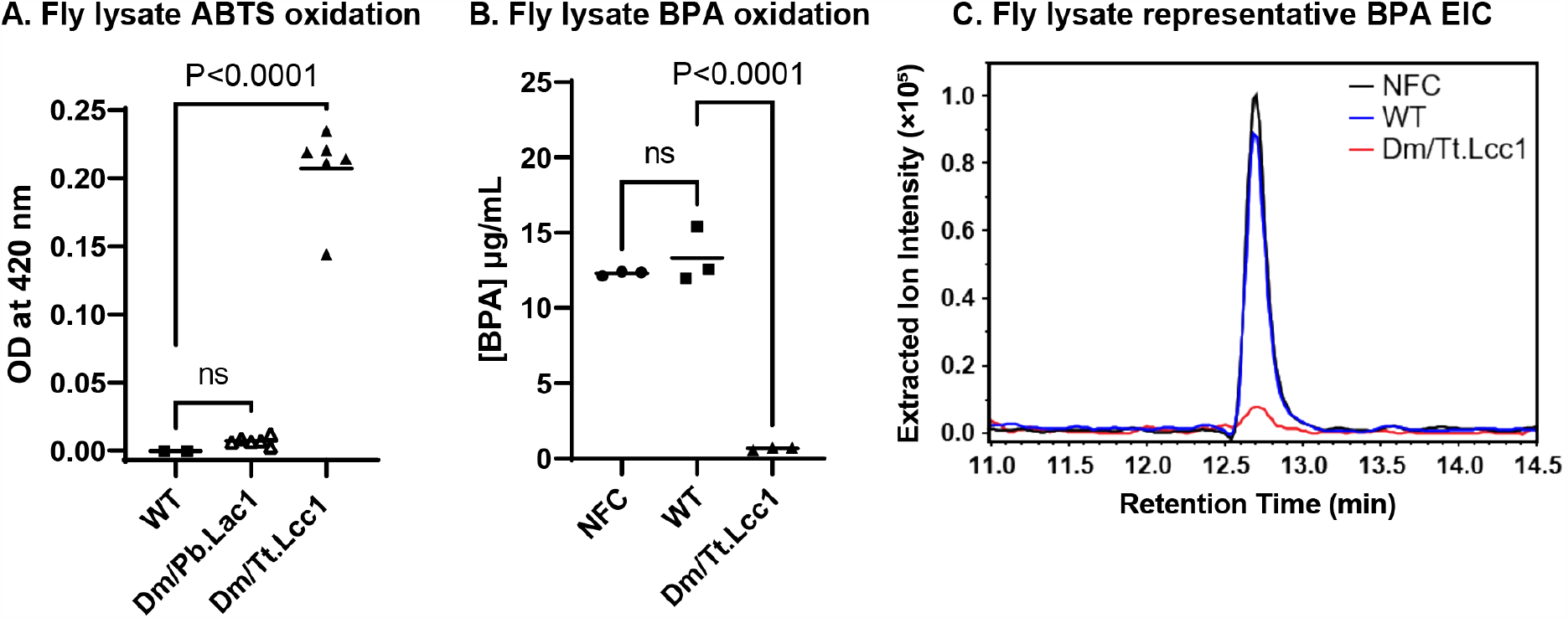
*In vitro* laccase substrate oxidation in fly lysates. **(A)** Fly lysate oxidation of 1 mg/mL ABTS for 2 hours. *n* = 2 biologically independent replicates for WT, *n* = 6 for all others. **(B)** Fly lysate degradation of 25 µg/mL BPA for 21 h. *n* = 3 biologically independent replicates. **(C)** Overlaid representative extracted ion chromatograms (EIC) (m/z 227.15) displaying the BPA peak for samples shown in **B**. Dm/Tt.Lcc1 = *D. melanogaster* engineered with *T. trogii* laccase, Dm/Pb.Lac1 = *D. melanogaster* engineered with *P. brumalis* laccase, WT = wild-type, NFC = No-Fly control. The statistical analysis was conducted using one-way ANOVA using Dunnett’s method compared to WT, where ns = not significant.

We next determined if Dm/Tt.Lcc1 flies are capable of degrading bisphenol A (BPA), an endocrine disruptor and emerging environmental contaminant found in many consumer products^47^. BPA has been shown to be a substrate of *T. trogii* laccase when expressed from its native host^48^. We incubated lysates from flies with 25 µg/mL BPA for 21 hours and measured BPA concentrations by liquid-chromatography mass spectrometry (LC-MS). Lysates from Dm/Tt.Lcc1 reduced BPA concentrations by 95% compared to the no-fly controls and the wild-type fly lysate (Fig. 2b-c).

### *In vivo* laccase activity

We evaluated the potential of laccase expressing insects to oxidize substrates present in their environment by secreted enzymes. Flies were reared on a minimal media to avoid binding of substrates to insoluble components present in standard media. The minimal media included 1 mM CuSO_4_ to supply the laccase co-factor. First, we reared 3 adult males and 3 adult females on minimal diet including 1 mg/mL ABTS for 2 days, after which they were removed. Larvae of the mated flies were grown for an additional 5 days. Dm/Tt.Lcc1 flies oxidized ABTS as indicated by blue colorization of the fly media, however, no oxidation was observed with wild-type flies (Fig. 3a). We controlled for the possibility that the ABTS oxidation was due to laccase that may have been leaking from dead larvae by incubating 6 male flies in ABTS media. After 11 days, only the Dm/Tt.Lcc1 males had oxidized ABTS (Fig. 3b). This result indicates that the oxidation was due to enzyme secreted into the media.

**Fig. 3:**
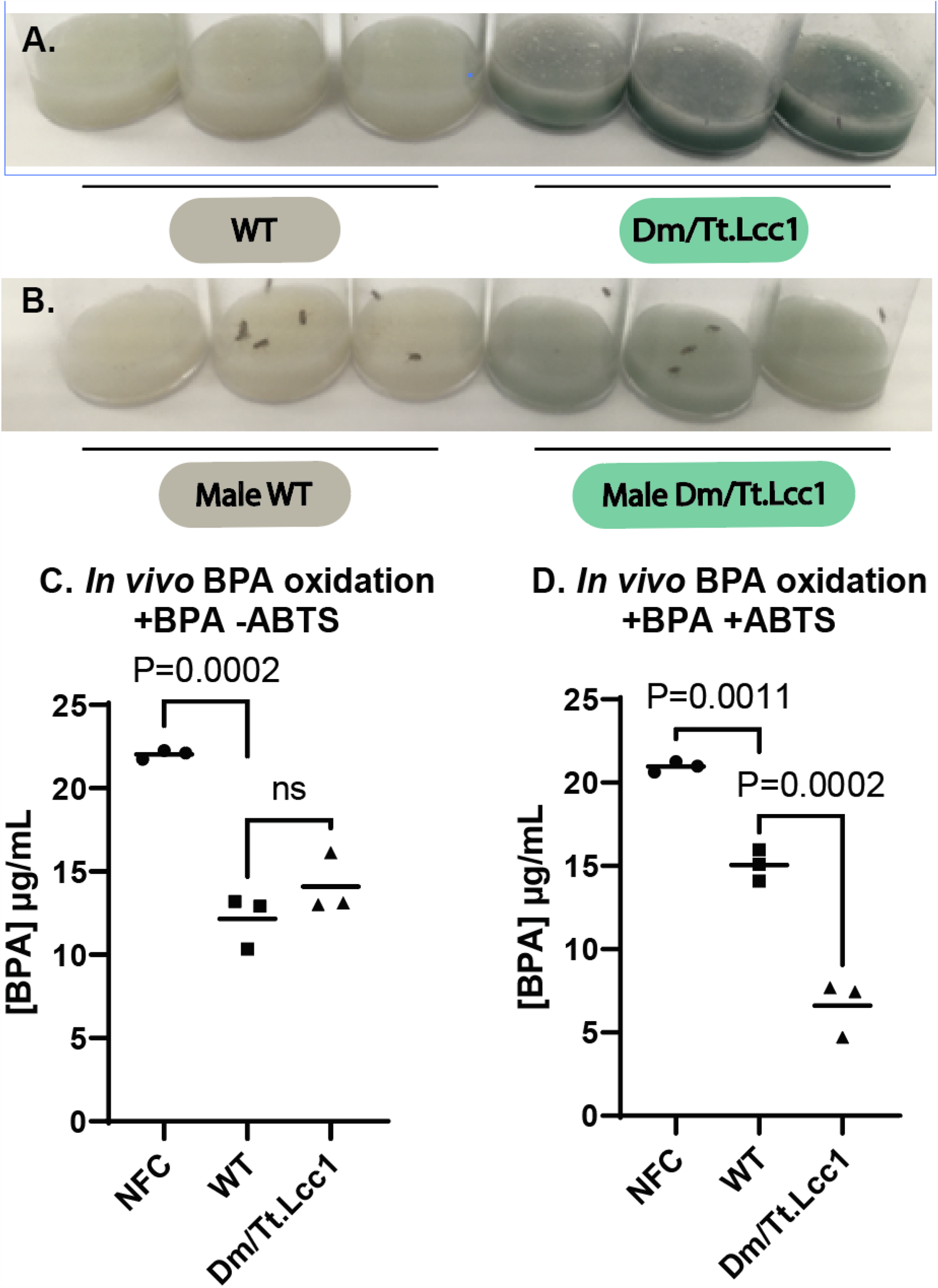
*In vivo* laccase enzyme activity. **(A)** *In vivo* oxidation of 1 mg/mL ABTS after 7 days. (**B)** *In vivo* oxidation of 1 mg/mL ABTS after 11 days using only male flies. **(C, D)** *In vivo* degradation of 25 µg/mL BPA without ABTS **(C)** and with 1 mg/mL ABTS **(D)**. *n* = 3 biologically independent replicates. Dm/Tt.Lcc1 = *D. melanogaster* expressing *T. trogii* laccase, WT = wild-type, NFC = No-Fly control. The statistical analysis was conducted using one-way ANOVA using Dunnett’s method compared to WT, where ns = not significant.

Next, we reared 5 male and 5 female flies in a minimal diet containing 25 µg/mL BPA. Adults were removed after three days. On day 10, BPA was extracted from their diet and quantified by Ultra-High-Performance Liquid Chromatography coupled with fluorescence detection (UHPLC-FLD). Both Dm/Tt.Lcc1 and wild-type flies had reduced BPA concentrations by 36% and 45%, respectively, compared to no-fly controls (Fig. 3c). There was no significant difference between Dm/Tt.Lcc1 and wild-type indicating that the reduction in BPA concentration was not due to the activity of the laccase transgene. The reduction in BPA concentration for wild-type flies relative to the no-fly control may be in part due to BPA absorbed by the adult flies which were removed after three days. Another possibility is that BPA detoxification pathways have been characterized in mammals^54^ and it is possible *D. melanogaster* also contains endogenous pathways capable of detoxifying BPA.

Some laccase substrates, such as ABTS, can function as redox mediators to facilitate the oxidation of additional compounds^55^. When *in vivo* BPA degradation was tested in the presence of ABTS, Dm/Tt.Lcc1 flies effectively reduced BPA concentration by 68% and 56% compared to the no-fly control and wild-type, respectively (Fig. 3d). It is not clear why ABTS was necessary for the *in vivo* assay and not the *in vitro* lysate assays. Adding lysate directly to the media also required ABTS for efficient degredation, suggesting that there is some component of the diet which may be interfering (Fig. S1).

### Lyophilized insect powder laccase activity

We next examined the potential of insects as platforms to produce a lyophilized laccase which can be easily stored and transported before use. A crude lyophilized powder from Dm/Tt.Lcc1 was compared to a commercially available purified laccase from *Trametes versicolor* using ABTS as a substrate. The lyophilized Dm/Tt.Lcc1 displayed a total laccase activity of 1.4 U/g ± 0.1 SEM dry weight, while the commercial purified laccase displayed a specific activity of 75.3 U/g ± 6 SEM dry weight.

We next tested if Dm/Tt.Lcc1 lyophilized whole-fly powder had activity against indigo carmine, a textile industry dye and pollutant^43^. Incubating 5 mg/mL of Dm/Tt.Lcc1 lyophilized powder in water containing 100 mg/L indigo carmine resulted in 90% decolorization after 48 h; however, WT fly lyophilized powder also decolorized 55% of the dye (Fig. 4a). We observed that insoluble material in the powder appeared to adsorb the dye. However, incubating dye using supernatant from centrifuged lyophilized Dm/Tt.Lcc1 in water did not appear to decolorize indigo carmine compared to controls (Fig. 4b). When the Cu^2+^ co-factor was included, supernatant from centrifuged lyophilized powder in water decolorized nearly 100% of indigo carmine in Dm/Tt.Lcc1 and 25% from WT flies after 90 hours (Fig. 4c-d).

**Fig. 4:**
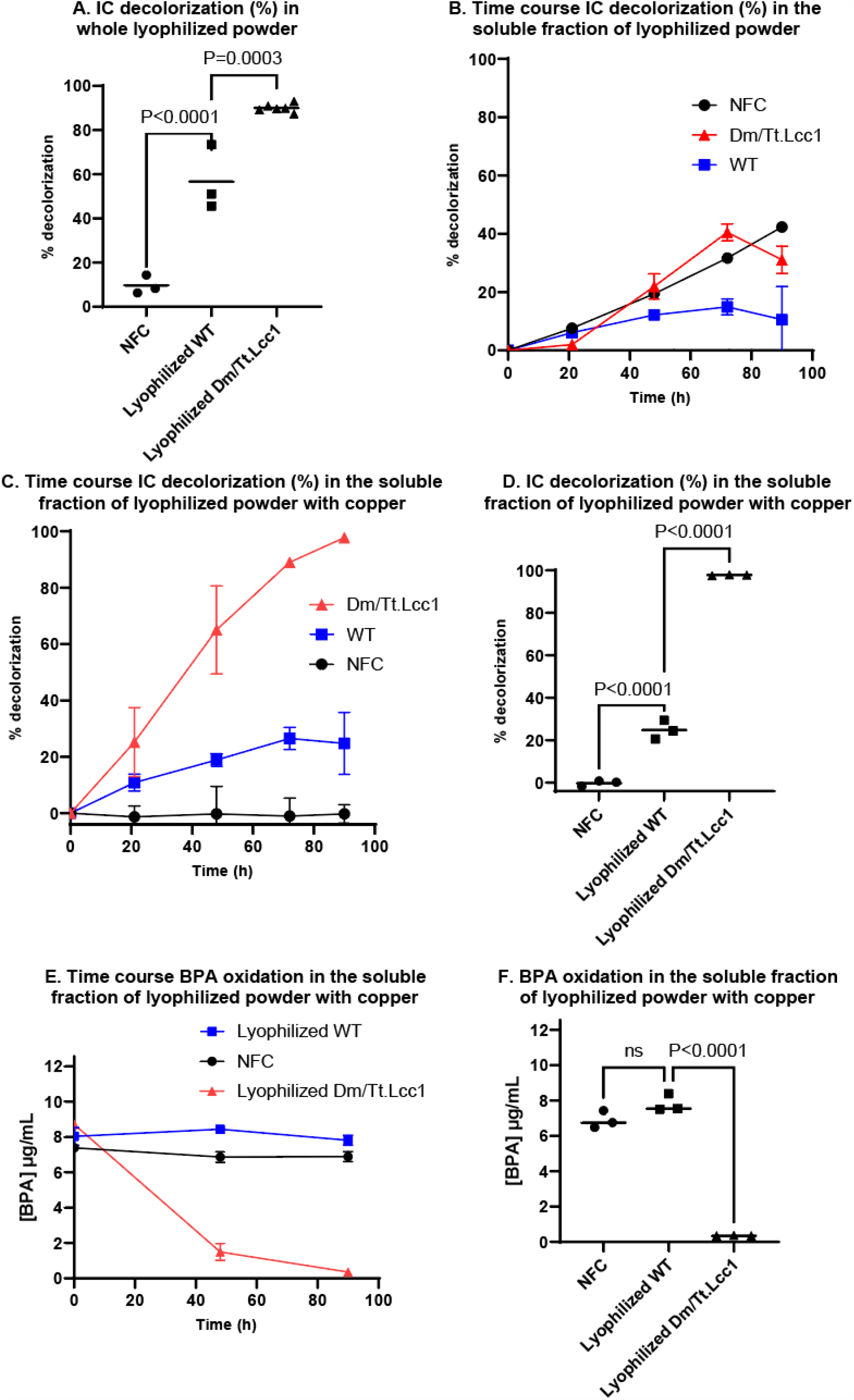
*In vitro* activity of lyophilized Dm/Tt.Lcc1 whole-fly powder on laccase substrates. **(A)** Decolorization (%) of 100mg/L indigo carmine at 610 nm in whole lyophilized powder resuspended in water after 48 hours. *n* = 3 biologically independent replicates. **(B-D)** Time course decolorization (%) of 100 mg/mL indigo carmine in the soluble fraction of lyophilized powder in water **(B)** or supplemented with 10 mM CuSO_4_ **(C)**. Statistical analysis of **C** was performed at end point **(D)**. *n* = 3 biologically independent replicates. **(E**,**F)** Time course degradation of 12.5 µg/mL BPA in the soluble fraction of lyophilized samples in water supplemented with 10 mM CuSO_4_ **(E)**. Statistical analysis of **E** was performed at the end point **(F)**. *n* = 3 biologically independent replicates. Dm/Tt.Lcc1 = *D. melanogaster* engineered with *T. trogii* laccase, WT = wild-type, NFC = No-Fly control. The statistical analysis was conducted using one-way ANOVA using Dunnett’s method compared to WT, where ns = not significant.

We also evaluated Dm/Tt.Lcc1 lyophilized powder activity against BPA. Incubating 12.5 µg/mL BPA with soluble enzyme from the lyophilized powder in the presence of Cu^2+^ co-factor resulted in a 96% reduction of BPA from Dm/Tt.Lcc1 fly powder after 90 hours. Incubation with WT fly powder resulted in a 3% reduction of BPA (Fig. 4e-f).

## Discussion

Engineering animals to heterologously express fungal/microbial enzymes may improve sustainable waste management, bioremediation, and facilitate the use of low-value waste streams as inputs for the production of high-value products such as industrial enzymes. Here, we engineer *D. melanogaster* to express a functional laccase from the fungus *T. trogii*. Lysates of this engineered fly (Dm/Tt.Lcc1) oxidized the endocrine disruptor BPA by 95% *in vitro* compared to the WT flies. Engineered flies also degraded BPA by 56% *in vivo* when added to their diet. Further, a powder prepared from lyophilized engineered flies also degraded BPA by 96% and the textile dye indigo carmine by almost 100% *in vitro*.

Mediator compounds are often used to improve the substrate range and activity of laccases^56^. Although Dm/Tt.Lcc1 lyophilized powder did not require the addition of ABTS as a mediator for degrading BPA, *in vivo* BPA degradation did. White rot fungi such as *Phanerochaete chrysosporium* and *Pycnoporus cinnabarinus* synthesize and secrete endogenous redox mediators^57,58^ and it may be possible to engineer animals to secrete redox mediators to also enhance their bioremediation activities. Alternatively, colloidal lignin or derivatives of lignin also show potential to act as redox mediators^59,60^, and including them may be feasible to assist with *in vivo* activity.

Using transgenic insects as platforms to convert waste to high-value products, such as industrial enzymes, will require high production titres. Dm/Tt.Lcc1 lyophilized powder made from whole flies had 1.4 U/g of laccase activity without any attempt at optimization compared to 75.3 U/g of activity from a commercially available purified *T. versicolor* laccase. This is a very promising start and developing tools to genetically manipulate more relevant hosts and strain engineering is needed for insects to reach their potential as bioproduction platforms.

In summary, our findings demonstrate for the first time that insects can be stably engineered to express a functional fungal laccase, which demonstrated *in vivo and in vitro* laccase activity. Transgenic flies were capable of oxidizing laccase substrates in their environment and they can be used as a lyophilized powder to oxidize substrates in aqueous solutions. These results provide strong support for further research in using transgenic animals as platforms for bioremediation, waste stream management, and recombinant protein production.

## Methods

### Construct design and assembly

NEBuilder® HiFi DNA Assembly Master Mix (New England Biolabs) was used to assemble all constructs. pMBO2744^15^ was used as a backbone to generate the enzyme expression plasmids. A gBLOCK (IDT®) was synthesized that encodes a short alpha-tubulin promoter, a downstream SV40 polyadenylation sequence, and a NotI restriction site in between. This gBLOCK was installed into NotI linearized pMB02744 by HiFi assembly to produce pMC-1-1-1 (Supplementary Table 4). This plasmid was then linearized with NotI, and gBLOCKs that encode for *T. trogii* laccase or *P. brumalis* laccase enzyme were installed with HiFi assembly to generate pMC-1-2-3 and pMC-1-2-6, respectively (Supplementary Table 4). The DNA sequences were codon optimized for expression in *D. melanogaster* using the IDT® codon optimization tool. The native fungal signal peptides were replaced with the larval cuticle protein 9 signal peptide. The Kozak sequence, AATCTTACAAA, was upstream of the start codon. See Supplementary Table 2 for plasmid descriptions and Supplementary Table 3 for primers used in this study.

### Insect rearing and generation of transgenic fly strains

Canton S wild-type flies were from the Bloomington Stock Centre (64349, RRID: BDSC 64349). Enzyme expression plasmids were purified using ZymoPURE™ II Plasmid Midiprep Kit (Zymo Research #D4200) and sent to BestGene Inc (Chino Hills, Ca) for φC31 mediated integration. All strains were maintained on a cornmeal diet based on the Bloomington standard Nutri-Fly formulation (catalog number 66-113; Genesee Scientific). In preparation for the *in vitro* or *in vivo* laccase enzyme activity assay, the diet for both the transgenic and wild-type strains was supplemented with CuSO_4_•5H_2_O (1 mM, Merck, CAS-no: 7758-99-9). Flies were reared in a controlled environment room at 25.0 °C, 75% humidity, and a 12 hour light/dark cycle with a 30 minute transition period. See Supplementary Table 1 for all of the fly strains used in this study. Biosafety approval was granted by the Institutional Biosafety Committee of Macquarie University (#5049).

### *In vitro* enzyme activity assay

To obtain a fly lysate, ten frozen adult flies were homogenized with a motorized pestle in the stated buffer supplemented with 10 mM CuSO4•5H2O and acid-washed glass beads. The samples were then agitated along a tube rack 10 times. Each tube was incubated for 15 minutes at 25 °C and centrifuged twice to remove insoluble debris.

For the ABTS oxidation assay, fly lysates were prepared in citrate-phosphate buffer (pH 5) and added to 1 mM ABTS. The tubes were incubated at 25 °C for 2.5 hours. The reaction was quenched by adding 5% trichloroacetic acid (TCA, Sigma Aldrich, catalogue #T0699, CAS-no:76-03-9). A UV-Vis spectrophotometer (Jasco V-760 UV-Vis spectrophotometer) was used to measure the absorbance at 420 nm. The measurements were blanked to an equivalent sample that did not contain ABTS.

For the BPA degradation assay, fly lysates or controls were prepared in sodium acetate buffer (pH 5) and added to 25 µg/mL BPA. The assay mixtures were incubated at room temperature for 21 hours. The reaction was quenched by adding 5% (v/v) trichloroacetic acid (Sigma Aldrich, catalog #T0699, CAS-no:76-03-9). BPA concentration was analyzed by LC-MS as described below.

### *In vivo* laccase activity assays

#### Minimal media

The minimal media components were for 100 mL: 1.58 g yeast, 0.66 g agar, 5 g sugar, 4 mL 1 M propionic acid (pH 4.4), 12.4 mg CuSO_4_•5H_2_O (1 mM final concentration). A buffer of 20 mM citrate-phosphate (pH 5) was added for the initial ABTS assays, 25 mM sodium acetate buffer (pH 5) was used for all BPA experiments.

#### ABTS only *in vivo* experiments

To minimal media 100 mg ABTS (1 mg/mL final concentration) was added. Three female and three male adult flies were incubated on the media for 48 hours, after which they were removed. For the all-male ABTS assay, the adult flies remained in the vial for 11 days. The appearance of the dark green radical cation was monitored and recorded with photographs.

#### For the ABTS + BPA experiments

To minimal media, BPA was added (25 µg/mL final concentration). ABTS (1 mg/mL final concentration) was added for the +BPA, + ABTS condition.

For each condition, the liquid media was pipetted into a 14 mL polypropylene round-bottom culture tube. 5 male and 5 female adult flies were incubated on the media for 3 days and then removed. For the lysate condition, 50 µL of lysate from ∼100 flies was added to the tube. After a total of 10 days the enzymatic reaction was stopped with TCA (5% v/v final concentration). The media was melted in a water bath, after which methanol (making ∼1:1 methanol: aqueous TCA for a two-fold dilution of BPA) was added, followed by vortexing.

The tubes were then incubated at 300 rpm and 37 °C for 30 minutes. After incubation, the tubes were centrifuged twice with the final supernatant reserved for BPA quantification by UHPLC-FLD as described below.

### Lyophilized Dm/Tt.Lcc1 enzyme activity assays

Frozen Dm/Tt.Lcc1 or wild-type flies were lyophilized (Alpha 1-4 LDplus, Christ) at −45 °C at mBar. The freeze-dried flies were homogenized in a porcelain mortar and pestle and stored at 4 °C.

For the activity assays, lyophilized Dm/Tt.Lcc1 and commercial *T. versicolor* laccase (Sigma-Aldrich, Catalogue 38429) were resuspended in 25 mM sodium acetate buffer (pH 5) supplemented with 10 mM CuSO_4_•5H_2_O. Lyophilized Dm/Tt.Lcc1 did not dissolve, and the soluble fraction was extracted at 25 °C for 15 minutes at 800 rpm and centrifuged twice to remove the insoluble debris. Enzyme serial dilutions were added in duplicate to 1 mM ABTS and ABTS oxidation was monitored on a PHERAStar FS plate reader (BMG Labtech) at 420 nm at 25 °C. One unit of enzyme activity was defined as the amount of enzyme that catalyzes the oxidation of 1 *μ*mol ABTS min^-1^ using the molar extinction coefficient 36 mM^-1^ cm^-1^ ^61^.

For the indigo carmine decolorization assay, 5 mg/mL of lyophilized powder or controls in Millipore water were added to 100 mg/L indigo carmine (Sigma-Aldrich catalog 131164) and incubated at RT. At 0 and 48 hours, indigo carmine decolorization was monitored on a PHERAStar FS plate reader (BMG Labtech) at 610 nm.

As indigo carmine appeared to adsorb to the lyophilized powder, the soluble fraction was tested for enzymatic activity. 5 mg/mL lyophilized powder or controls in Millipore water supplemented with and without 10 mM CuSO_4_•5H_2_O were extracted at 25 °C for 15 minutes at 800 rpm. The samples were centrifuged twice to remove the insoluble debris. The supernatant of the lyophilized powder and controls were added to 100 mg/L indigo carmine and incubated at room temperature. Indigo carmine decolorization was monitored on a PHERAStar FS plate reader (BMG Labtech) at 610 nm at 0, 21, 48, 72, and 90 hours.

For the BPA degradation assay, the soluble fraction of the lyophilized powder was extracted as described in the indigo carmine assay. The supernatant of the lyophilized powder and controls were added to 12.5 µg/mL BPA and incubated at room temperature. At 0, 48, and 96 hours, 100 μL aliquots were removed and the reaction was stopped using a 3 KDa nominal molecular weight limit (NMWL) centrifuge filter (Merck Amicon Ultra 0.5mL Centrifugal Filters, UFC500396). BPA concentration was analyzed by UHPLC-FLD as described below.

### BPA quantification

BPA in samples was either quantified by UHPLC-FLD analysis or by LC-MS. UHPLC-FLD analysis was performed using an Agilent RRHD ZORBAX Eclipse Plus C18 column (3.0 × 150 mm 1.8 μm, p/n 959759-302) equipped with a UHPLC Eclipse Plus C18 Guard Column, (2.1 mm 1.8 µm, p/n 821725-901). The isocratic mobile phase consisted of acetonitrile (ACN):10 mM KH2PO4 (50:50 v/v) at a flow rate of 0.5 mL/min. A gradient wash step was introduced after each run with eluent B 100% ACN and eluent A 10mM KH2PO4, starting with 45% B1 from 0 - 1 minute, 70% B from 1 – 6 minutes, 5% B from 6 – 11 minutes, and finally 50% B from 11 - 16 minutes at 0.3 mL/min. BPA was detected by setting FLD excitation wavelength at 229 nm, and emission wavelength at 316 nm and a gain setting of 12. Linear regression of 7-point standards from 12.5 ng/mL to 12,500 ng/mL BPA was performed to calibrate BPA concentration. All samples and standards were either filtered using syringe filters (Filtropur S µm, Sarstedt, 83.1826.001) or centrifuge filters (Merck Amicon Ultra 0.5mL Centrifugal Filters, UFC500396) prior to injection into the UHPLC.

LC-MS validation and quantification of BPA was performed externally at the Mass Spectrometry Facility at the University of Sydney. LC separation was performed using a Waters Sunfire C18 column (5 *μ*m, 2.1 mm ID x 15 mm) coupled to a Bruker amaZon SL mass spectrometer. Gradient elution was performed with a mobile phase consisting of (A) methanol and (B) water with 0.1% acetic acid at a flow rate of 0.3 mL/min. The gradient consisted of 20% A from 0 - 9.3 minutes, 76% A from 9.3 - 17.3 minutes, 92% A from 17.3 – 26 minutes, and finally 20% A from 26 - 35 minutes. MS analysis was performed using Atmospheric Pressure Chemical Ionisation (ACPI) under positive and negative ionization mode over a range of 200-1400 m/z. The operation parameters were: Nebulizer 27.3 psi, Dry Gas 4.0 L/min, Dry temperature 180 °C, APCI Vaporisation temperature: 400 °C, APCI Corona needle current: 6000 nA, Mass mode: enhanced resolution, SPS (Smart parameter setting) Target mass 150 m/z, compound stability 100%. Linear regression of 6-point BPA standards from 20 ng/mL to 10 µg/mL BPA was performed to calibrate BPA concentration. An internal deuterated BPA-D_16_ standard was spiked into the samples prior to injection to calibrate ionization in the sample matrix compared to the standard curve matrix prepared in methanol. All samples were centrifuge filtered (Merck Amicon Ultra 0.5mL Centrifugal Filters, UFC500396) prior to injection into the LC-MS.

## Statistical analysis

Figures were generated in Prism 9 by GraphPad. For statistical analysis, a one-way ANOVA using Dunnet’s method was conducted. The significance level for the analysis was 0.05, and all error bars represent the standard error of the mean. The exact *P* values, statistical tests used, and sample numbers (*n*) can be found in the figure legend.

## Acknowledgements

We are grateful Dr. Nick Proschogo from the Mass Spectrometry Facility at the University of Sydney.

## Author Contributions

MC, KT, and MM conceived of this study. MC, KT, KP, AS, and MM designed experiments and analyzed data. MC, KT, KP, and SK performed the experiments. All authors contributed to writing the manuscript.

## Data and materials availability

All data is available upon request.

## Supplementary Materials

Supplementary Figure 1

Supplementary Tables 1-4

Supplementary Data File

## Supplementary information

**Supplementary figure 1:**
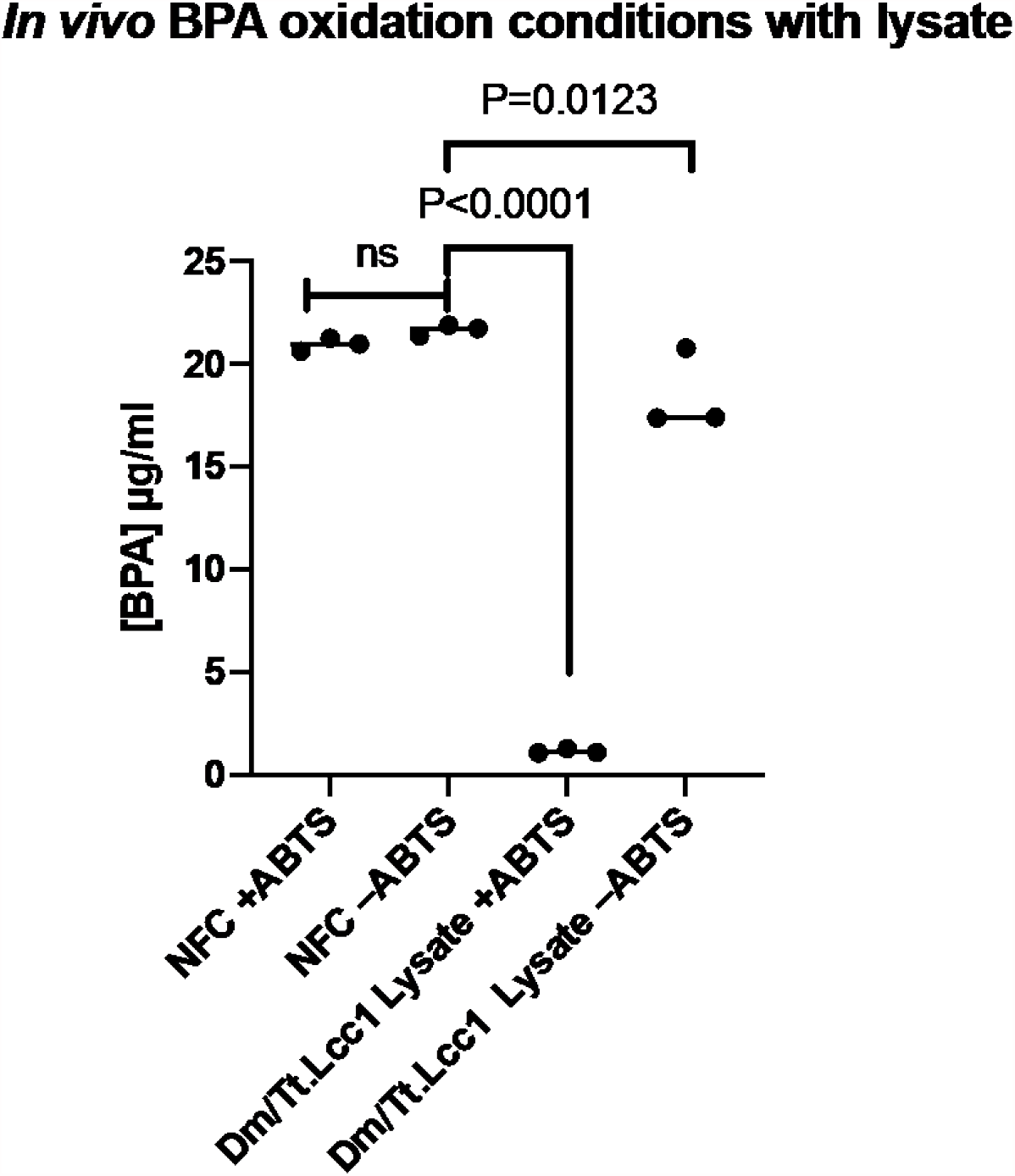
*In vivo* BPA degradation conditions comparing Dm/Tt.Lcc1 lysate to NFC. Fly lysate was added to media that contained 25 µg/mL BPA with and without 1 mg/mL ABTS. Dm/Tt.Lcc1 lysate was unable to degrade BPA without ABTS. *n* = 3 biologically independent replicates. The statistical analysis was a one-way ANOVA with Dunnett’s method. ns= not significant, Dm/Tt.Lcc1 = *D. melanogaster* engineered with *T. trogii* laccase, NFC = no fly control.

**Supplementary Table 1:**
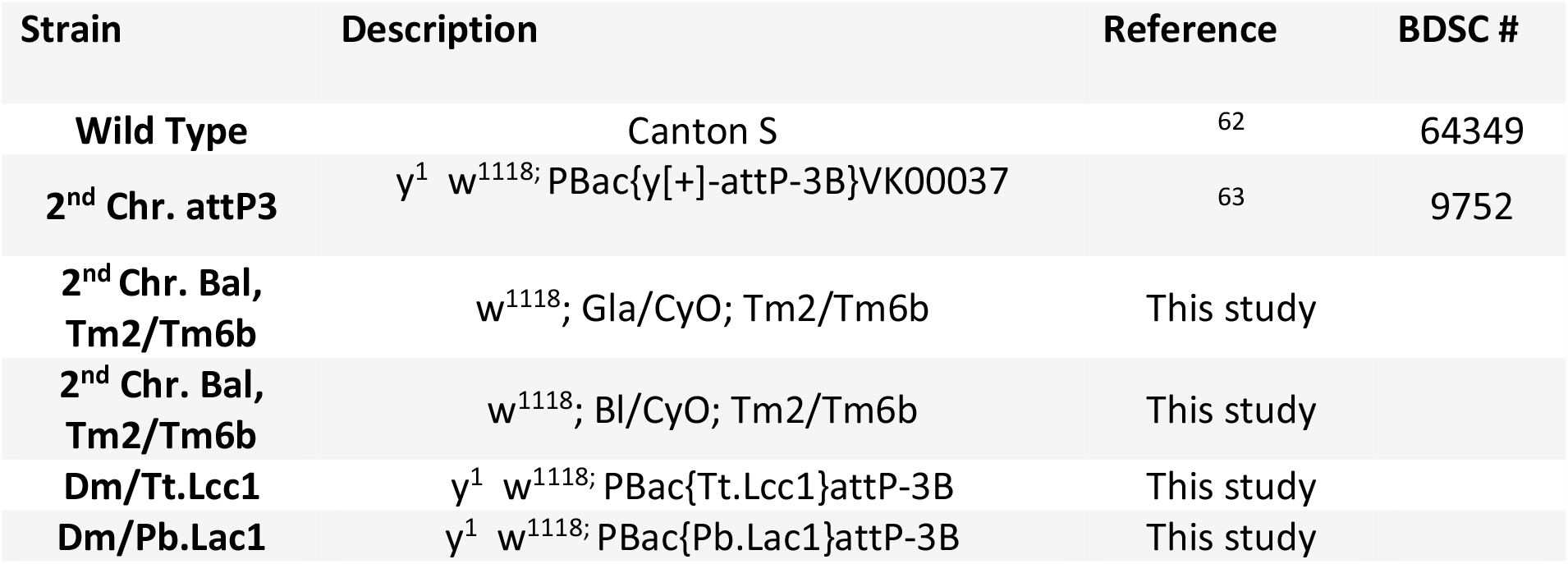
*D. melanogaster* strains used in this study.

**Supplementary Table 2:** Plasmids used in this study.

| Plasmid | Integrated in Fly Strain | Description | Reference/ GenBank Accession # |
| --- | --- | --- | --- |
| pMB02744 | NA | Contains a mini-white gene for *D. melanogaster* selection, an *AttB* site for genomic integration, and an ampicillin resistance cassette for cloning in *E. coli*. | 15 |
| pMC-1-1-1 | NA | Empty vector, with short alpha-tubulin promoter and SV40 regulatory sequences | This study |
| pMC-1-2-3 | Dm/Tt.Lcc1 | Short alpha-tubulin promoter driving expression of *Lcc1* from *Trametes trogii* | This study |
| pMC-1-2-6 | Dm/Pb.Lac1 | Short alpha-tubulin promoter driving expression of *Lac1* from *Polyporus brumalis* | This study |

**Supplementary table 3:**
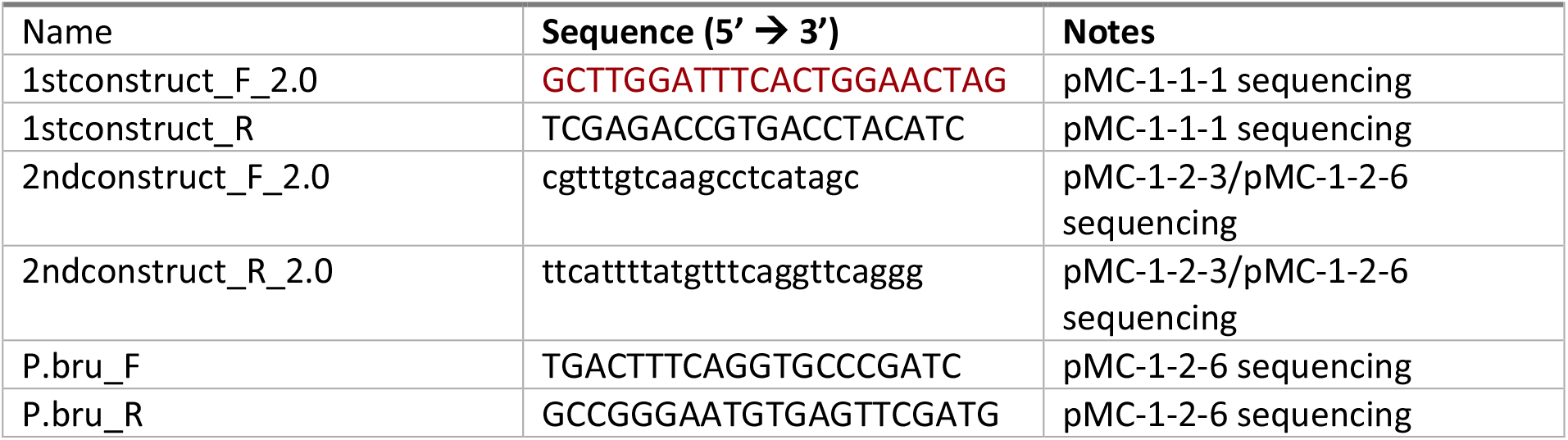
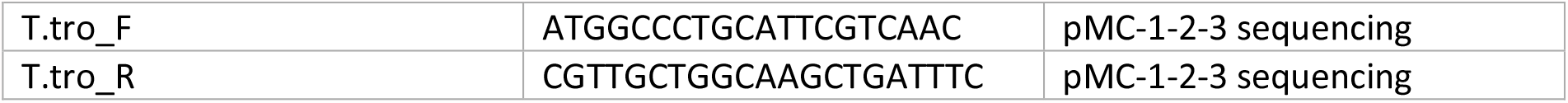
Primers used in this study.

**Supplementary table 4:**
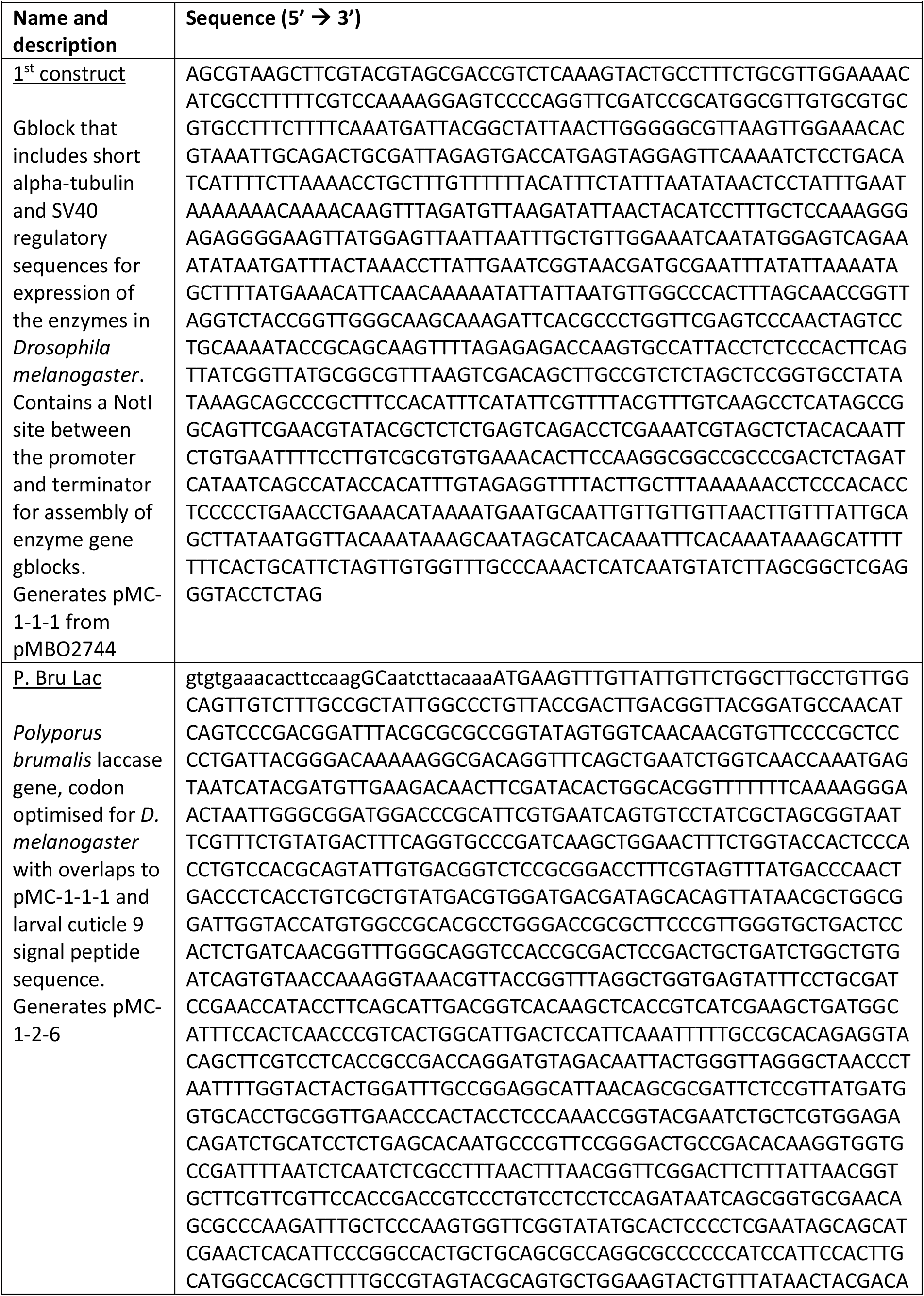
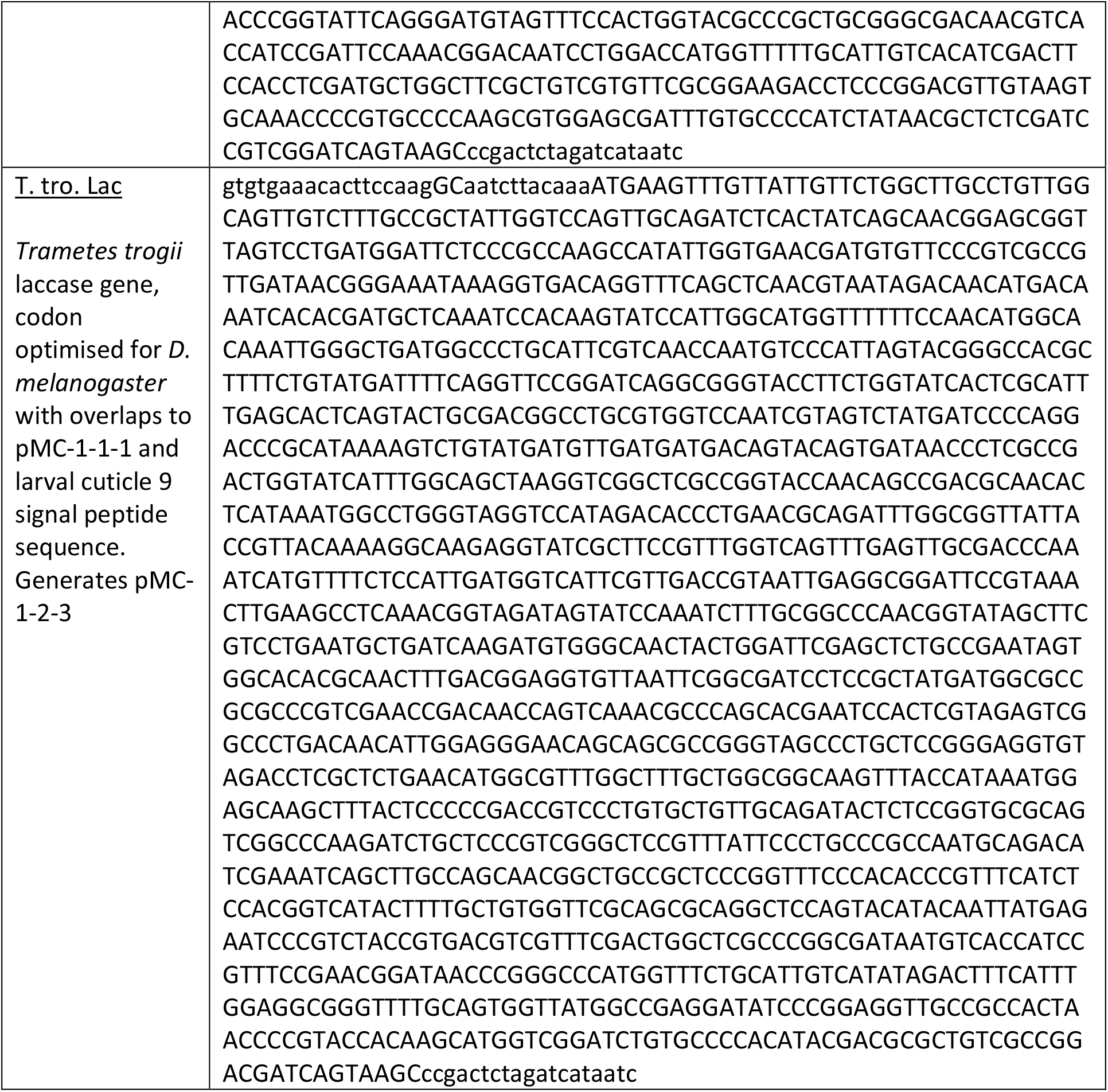
gblocks used to construct laccase expression plasmids:

